# Impact of databases on genomic survey of *Salmonella* antimicrobial resistance and virulence factor functional genes across the African continent

**DOI:** 10.64898/2026.01.14.699574

**Authors:** Balasubramanian Ganesan, Hana Azuma, Robert C. Baker, Jorge Pinto Ferreira, Jeffrey T LeJeune

## Abstract

Tracking the antimicrobial resistance potential of foodborne bacteria remains relevant today due to the potential for functional availability and horizontal transfer of relevant genes across species. We conducted a study of AMR potential across 4,552 *Salmonella enterica* isolates submitted to the NCBI Sequence Read Archive from the African continent. After assembling raw fastq data to high quality and contiguity, AMR genes, virulence factors, and plasmids and other mobile elements were predicted with the ABRicate software suite across multiple AMR databases. Prevalence of AMR genes in countries did not correlate with the number of *Salmonella* isolates sequenced. Isolates carrying functional genes were also classified by FAO food categories, identifying land animals meat and poultry and dairy as the primary sources of AMR genes amongst *Salmonella* isolates in Africa. The trends of AMR presence varied across different databases; whereas, the overall AMR genes per isolate for the same database did not vary substantially. Genomic comparison by k-mer analysis suggests most sampled isolates are not closely related within or across countries. Due to the low likelihood of cross border transmission, epidemiological reasons for AMR transmission within *Salmonella* in the five countries with highest prevalence of AMR genes are not identifiable presently.

## 1. Introduction

Across the last decade, whole genome sequencing (WGS) of microbial isolates, particularly of foodborne microbes, has grown incrementally. Efforts in this space were catalyzed by the 100k foodborne pathogen sequencing project [1], and subsequently, by increasing use of microbial WGS by national regulatory agencies such as the US Food and Drug Administration (FDA) [2], US Center for Disease Control (CDC) [2], The European Sood Safety Authority, Public Health England [3], and other national regulatory agencies. These efforts have been pivotal in creating the pathogens database at NCBI (https://www.ncbi.nlm.nih.gov/pathogens/) to provide the data from such efforts globally for research and development and technical validation. The United States National Antimicrobial Resistance Monitoring Systems (NARMS, https://www.cdc.gov/narms/index.html) currently uses genomic data as input for identifying antimicrobial resistance (AMR) potential and in the surveillance of genetic capabilities for AMR across the various collected isolates.

In this context, as more microbial isolates continue to be sequenced, the knowledge base of genetic and functional potential of microbes continues to evolve exponentially [4]. AMR is one such potential that is regularly tracked by NARMS and other national efforts (amrcountryprogress.org, https://www.who.int/activities/monitoring-progress-antimicrobial-resistance). The immediacy of resolving ever expanding AMR capabilities in difficult to treat pathogens is well established globally. Awareness of AMR issues is growing in low- and middle-income settings [5,6]. Guidelines for antimicrobial susceptibility monitoring and molecular testing are also available from some agencies, including Codex [7] and NARMS; whereas, lack of consistent adoption of responsible and prudent antimicrobial use in human medicine and agricultural production continues to contribute to the increase in microbial AMR potential [8].

Historically, AMR increase has been rarely attributed to foodborne pathogen linked transmission. This is primarily due to the lack of knowledge of AMR functional genes in foodborne pathogens [9,10]. Additionally, traditional testing for resistance across a broad spectrum of antimicrobials for determining minimal inhibitory concentrations is not always practicable in all nations. At many locations, the antibiotic itself may not be even available for therapeutic uses, and thus, even less available for diagnostic purposes [11],.

The application of whole genome sequencing may play a critical role in overcoming the need of antibiotics for AMR testing. Increasing adoption of whole genome sequencing of microbial isolates (www.ncbi.nlm.nih.gov/pathogens) contributes to the ability to mine functional genes contributing to AMR. The genome of an isolate at Africa or USA or Asia may be easily compared to large databases of AMR genes that are actively curated to a high quality. Many such databases are available for example, CARD [12], ResFinder [13], and AMRFinderPlus [14] are all bioinformatic tools to identify AMR genes supported by well curated databases. However, the number of genes curated and their variants are different across different databases (Table 1). This makes choosing and querying microbial genomic data against these databases to clearly identify patterns of AMR gene transfer challenging, and limits use of the results in a globally actionable manner.

**Table 1.** Isolate count per country.

| Country | Isolates per country |
| --- | --- |
| Algeria | 25 |
| Angola | 3 |
| Benin | 51 |
| Burkina Faso | 23 |
| Cameroon | 61 |
| Cape Verde | 2 |
| Central African Republic | 18 |
| Chad | 3 |
| Comoros | 6 |
| Djibouti | 2 |
| Democratic Republic of Congo | 130 |
| Egypt | 88 |
| Ethiopia | 375 |
| Gabón | 1 |
| Gambia | 167 |
| Ghana | 77 |
| Kenya | 163 |
| Liberia | 2 |
| Madagascar | 22 |
| Malawi | 349 |
| Mali | 360 |
| Mauritania | 7 |
| Mauritius | 14 |
| Mayotte | 2 |
| Morocco | 50 |
| Mozambique | 4 |
| Namibia | 6 |
| Niger | 6 |
| Nigeria | 609 |
| Réunion | 20 |
| Rwanda | 32 |
| Senegal | 157 |
| Sierra Leone | 6 |
| South Africa | 1326 |
| Sudan | 9 |
| United Republic of Tanzania | 119 |
| Togo | 16 |
| Tunisia | 56 |
| Uganda | 80 |
| Zambia | 78 |
| Zimbabwe | 31 |

The impact of AMR varies in different continents. Regions with poor Water, Sanitation and Hygiene (WASH) infrastructures, regions with inefficient infection, prevention and control procedures, regions where regulations on antimicrobial use are poorly enforced, and regions with developing healthcare systems and/or high levels of poverty tend to be more severely impacted. For example, African countries in the sub-Saharan region had the highest mortality rate (23.5 deaths per 100,000) in 2019 attributed to AMR compared to other regions [15]. Multiple studies report high prevalence of resistant isolates in countries like South Africa [16] and Ethiopia [17], emphasizing the need for integrated One Health surveillance for AMR and antimicrobial use (AMU), and the deployment and application of innovative source attribution methods.

*Salmonella* species are widespread and an important cause of human foodborne infections in Africa, such as typhoid fever [18], caused by *Salmonella* Typhi and *Salmonella* Paratyphi. Other clinical syndromes include diarrheal disease are caused by many non-typhoidal *Salmonella* Serovars [16,17]. Monitoring the genetic gain and loss or even the broader prevalence of AMRs within *Salmonella* may provide an important indicator of foodborne AMR transmission. Such surveys however are sparse, even in the light of NARMS and other survey efforts, particularly for the African continent. Recently, the occurrence and transmission of invasive non-typhoidal *Salmonella* has been characterized using whole genome sequencing for 1,300 isolates in sub-Saharan Africa [19]. With increasing isolate sampling and genetic diversity, the AMR gene content may also vary in different ways. However, performing such large scale genomic and genetic survey is additionally complicated due to the lack of standardization of gene content in AMR gene databases that may possibly lead to variable interpretations of the output.

In this study, we have attempted to assess this gap using available genomic data in INSDC databases for *Salmonella* spp. in various African countries, as of June 2021. We hypothesized that AMR gene content in *Salmonella* across African countries is not correlated to number of available isolates per country. We acquired unassembled fastq data from INSDC, trimmed, and assembled them into high quality draft genomes. These genomes were compared against various AMR gene databases in order to compile a survey of AMR and other functional genes that helps us understand factors such as genomic diversity, in relation to sampling density, and geographic distribution within the African continent.

## 2. Materials and Methods

### 2.1. Genome raw sequence data sources

Whole genome sequencing data of *Salmonella* was obtained as fastq files from the NCBI SRA database (www.ncbi.nlm.nih.gov/sra; one arm of INSDC), selected for only isolates that were sequenced on the Illumina platform in paired format only. Isolates from other platforms were fewer than 20 and were not included to maintain platform consistency for comparisons.

Country specific isolates were obtained by searching by each individual country’s name, for example, “*Salmonella* AND Algeria”, for all 55 African countries, dependencies/departments and territories. The total counts of isolates per country is provided in Table 1. Sequencing data was available only for the 42 countries, dependencies/departments and territories listed in Table 1. No sequencing data was available for following African countries, dependencies/departments and territories: Botswana, Burundi, Cote d’Ivoire (Ivory Coast was also used as a search term), Republic of Congo, Equatorial Guinea, Eritrea, Eswatini, Guinea, Guinea Bissau, Lesotho, Libya, Sao Tome & Principe, and Seychelles.

All raw sequencing data was imported via NCBI SRA toolkit v 2.9 or higher using the prefetch and fastq-dump utilities. Raw fastq data was trimmed to remove sequencing adapters using Trimmomatic software v0.32 [20] with default settings. Subsequently, phiX vector sequence removal was performed with bowtie2 [21], the filtered reads were merged with FLASH [22], and then assembled using SPAdes v3.13.0 or higher [23].

### 2.2. Post assembly comparison

Upon completion of genome assembly, assembly quality was compared using QUAST software v4.6.0 [24]. High quality genome assemblies were defined as those with a minimum genome size of 4,000,000 bp, minimum N50 of 50,000, and with fewer than 200 contigs. Genomes were compared for diversity by genome distance using Mash software v2.0 [25] with k-mer size of 31 and sketch size of 100,000. Closely related isolates were determined as those with the least Mash distance value, whereas identical pairs of isolates determined as those with a genome distance of 0. We also used the shared k-mers output from Mash for determining isolate relatedness with two ranges: ≤ 10 shared k-mers (identified as “indistinguishable”), and ≥ 10 and ≤ 100 shared k-mers (“distantly related”). *Salmonella* serovar for each isolate was predicted using SISTR software v1.10 or higher [26].

AMR genes in each isolate assembly were predicted by the ABRicate software (release dt. Feb. 2020 or later; github.com/tseemann/abricate) against all supported pre-downloadable databases (NCBI AMRFinderPlus, CARD, ResFinder, ARG-ANNOT, MEGARES, EcOH, PlasmidFinder, VFDB, and Ecoli_VF). ABRicate depends on BLASTN v2.7 or higher [27] and any2fasta (v0.4.2, github.com/tseemann/any2fasta) software.

### 2.3. Statistics and data visualization

Across country counts for AMR genes identified were plotted in Microsoft Excel software. Statistical differences of AMR gene counts predicted for each database across countries were assessed pairwise by chi-square tests using Excel. Statistical significance was assigned at α = 0.05.

## 3. Results

We collated whole genome sequence data for the 42 African countries as described above. Across 4,552 high quality genomes, 2,433 were from humans, and 1002 from food sources, and the rest from unidentified sources. The raw sequence data was assembled into genome scaffolds, and these scaffolds were submitted as queries to various databases supported by the ABRicate software. We found 881,010 matches at the nucleotide level to various functional genes within these databases. The percent identity across the entire length of genes was extracted from ABRicate output for quality checking (Table 2). Of these matches, 75% were 100% identical and 99% were at least 90% identical to the known resistance or virulence factor or plasmid genes within these databases. Also, all matches except two were matching at ≥ 80% nucleotide level identity. As this is relatively stringent, we felt confident that the predictions are likely accurate and useful for broader statistical comparisons.

**Table 2.** Matches to AMR genes across multiple databases.

| Database | Isolates | Total | Per<br>isolate | Type | Gene<br>count | DATE |
| --- | --- | --- | --- | --- | --- | --- |
| argannot | 4552 | 25015 | 5.5 | AMR | 1749 | 2019-Jul-28 |
| card | 4556 | 124010 | 27.2 | AMR | 2241 | 2019-Jul-28 |
| ecoh | 62 | 126 | 2 | E. coli O-groups and H-types | 597 | 2019-Jul-28 |
| ecoli_vf | 4554 | 185540 | 40.7 | E. coli VFDB Virulence factors | 2701 | 2019-Jul-28 |
| megares | 4556 | 165544 | 36.3 | AMR | 6635 | 2020-Feb-20 |
| ncbi | 2011 | 11384 | 5.7 | NCBI AMRFinderPlus | 4324 | 2019-Jul-28 |
| plasmidfinder | 2863 | 6752 | 2.4 | Plasmids | 263 | 2019-Jul-28 |
| resfinder | 4528 | 15824 | 3.5 | AMR : acquired genes, chr mutations | 2434 | 2019-Jul-28 |
| vfdb | 4556 | 469042 | 103 | Virulence factors | 2597 | 2019-Jul-28 |

Initially, we compared the predicted counts of three groups of functional genes, i.e. AMR, virulence, and plasmid genes within each country (Fig. 1). For this purpose, the sum of all AMR, virulence, and plasmid genes predicted across all reference databases was compared, irrespective of potential duplicates across the databases. Thus, the overall numbers may be slightly inflated, however we don’t expect that the trends will significantly differ. The count of detected genes ranged between 150-334. Most countries had gene counts per isolate between 180-200, with five countries falling outside the first quartile (Fig. 1).

**Fig. 1.**
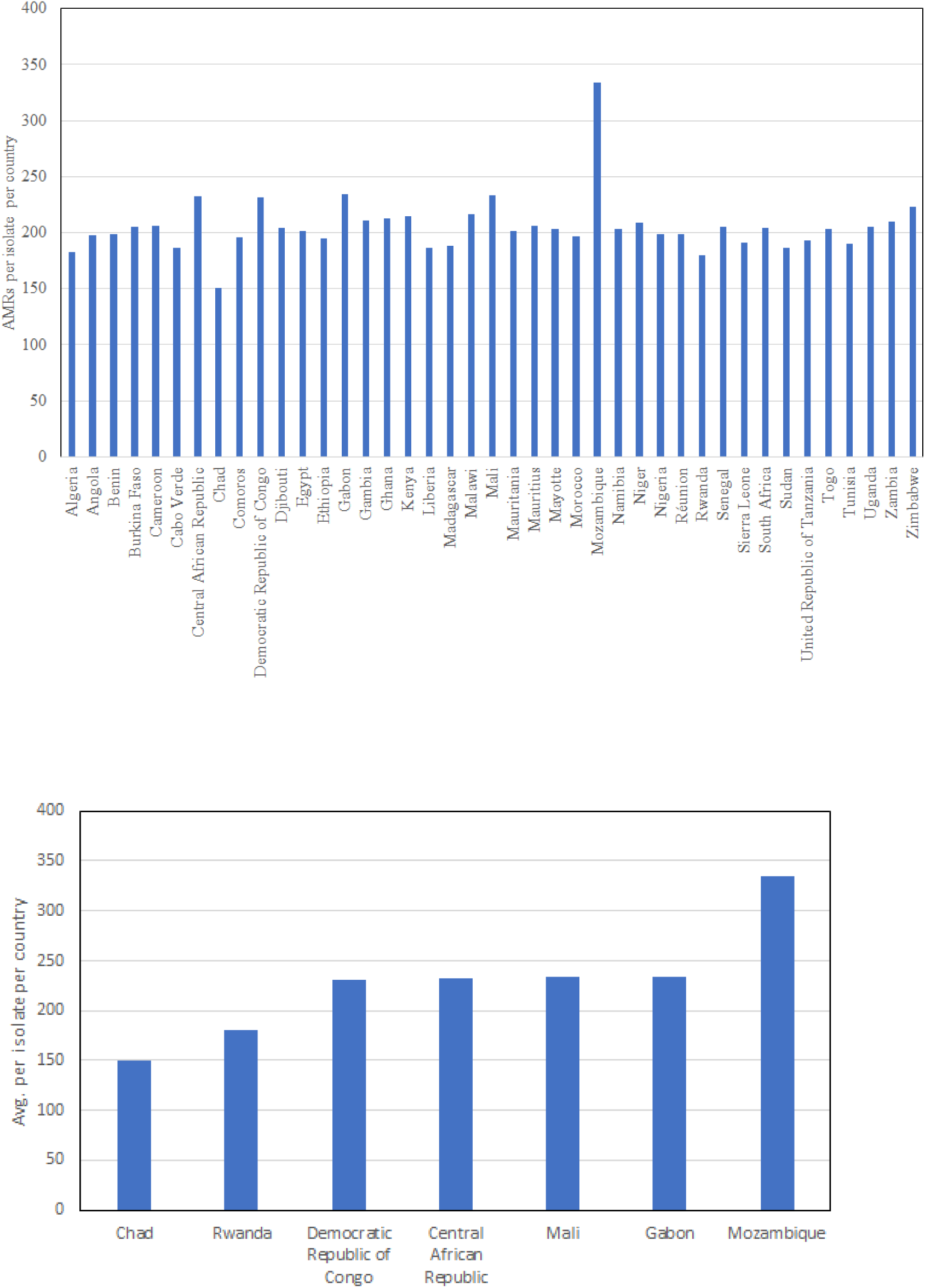
AMR count per isolate per country from 42 countries (upper panel) and countries outside the first quartile (bottom panel)

The ABRicate software supports nine different databases, five of which are directly collections of AMR genes (argannott, card, megares, NCBI AMRplus, and resfinder; Table 2). With increasing numbers of genes across databases, we expected that the number of AMR genes discovered would cumulatively be higher. However, we found no strong correlation across databases in number of AMRs found total per country or per isolate (Table 2). The average number of AMR genes was 5.5 for argannot, 27 for card, 36 for megares, 5.6 for NCBI AMRfinderplus, and 3.5 for resfinder databases. For these 5 databases, where matches were found, the least number of matches per isolate was 1, except for card with 7 matches minimum. Notably, all isolates had at least 1 hit within card and megares, whereas 4 isolates had no matches to argannot, 28 isolates had no matches to resfinder, and 2,545 isolates had no matches to NCBI AMRfinder plus database. The NCBI AMR finderplus and megares databases have the highest gene counts within, yet 55.8% isolates had no matches to the first database. The maximum number of matches for an AMR gene database for an individual isolate ranged from 20-91. The distribution of total AMR counts found across all countries (Fig. 2) were statistically compared across pairs of AMR databases using a chi-square test, and every pairwise comparison was statistically significant (P < 0.05), suggesting that the same results may not be obtained when different reference databases are chosen for AMR surveillance. These observations highlight the variability when specific AMR reference databases are chosen for surveillance.

**Figure 2.**
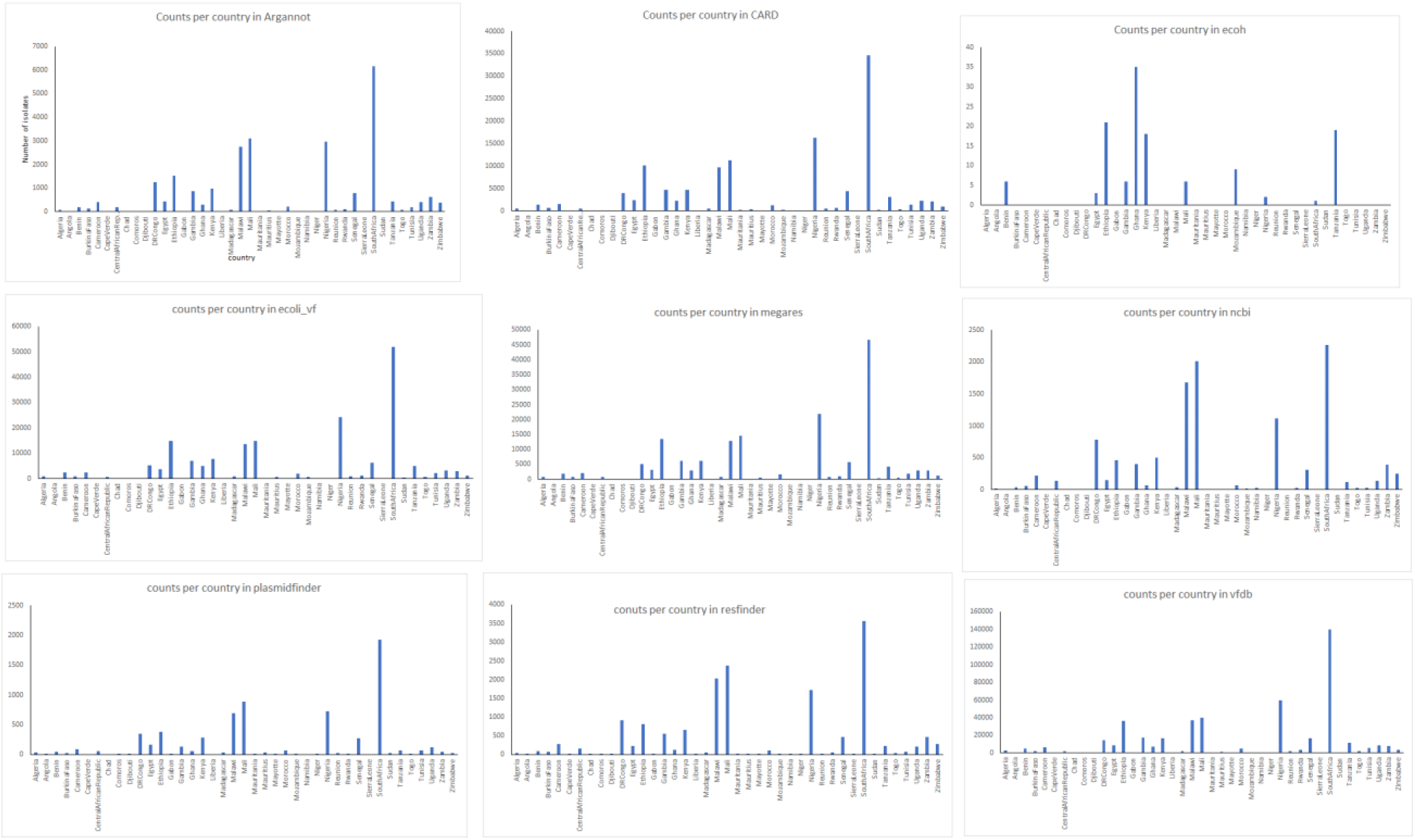
Country-wise distribution of various functional genes overall

Plasmids are known to carry AMR genes. Plasmid mobility is a key mechanism for transferable AMR capability. We assessed the prevalence of plasmid-borne genes by comparing the assembled 4,556 genomes against the plasmidfinder database. This was necessary as the genomes are in scaffold form and not completely circularized. Out of 4,556 assembled genomes, 2,863 (63%) carried plasmid-borne genes, whereas the remaining 37% had no matches to known plasmid-borne genes. However, apart from 3 countries that had no plasmid gene hits, Cape Verde, Chad, and Sierra Leone with 6, 6, and 18 isolates respectively, every other country’s isolates had at least a total of 2 and as many as 1,924 plasmid-borne gene matches in total. This highlights the broad prevalence of plasmids within *Salmonella* genomes from the African continent.

We assessed the prevalence of virulence related genes supported by matches to three databases supported by ABRicate (Table 3, Fig. 1). Matches to the ecoh databases were relatively low (only 62 of 4,556 isolates showing total 126 hits), which is to be expected due to *Salmonella* isolates being compared against an *E. coli*-specific database. However, almost every isolate had matches to the ecoli_vf and vfdb databases. This is to be expected by BLASTN homology as *Salmonella* and *E. coli* virulence factors may share high homology even at the genetic level. As the majority of isolates were sourced from clinical samples, it is to be expected that the isolates are most likely to carry virulence factor genes, which is confirmed now by assessing virulence factors by DNA sequence homology.

**Table 3.** Number and percentage of collective BLASTN matches across all seven functional gene databases at different ranges of identity.

| Identity (%) | Number of hits* | Percent in range |
| --- | --- | --- |
| 100 | 664,928 | 75.47 |
| < 100 | 216,082 | 24.53 |
| < 95 | 28,308 | 3.21 |
| < 90 | 8,888 | 1.01 |
| < 85 | 3,091 | 0.35 |
| < 80 | 2 | 0.00 |

\*Total matches for all isolates across all databases = 881,010

We compared the overall prevalence between high and low counts of total AMR, virulence, or plasmid genes. From a country-wise perspective, for overall functional genes, South Africa, Malawi, Mali, and Nigeria were the countries with highest gene counts in these categories across all databases except ecoh. However, the number of isolates and number of AMR, plasmid, or virulence genes across countries are linearly correlated (Fig. S1, supplementary data), indicating that as more isolates are sampled across countries, more genes related to these capabilities are likely to be detected.

To avoid sampling bias due to higher isolate counts, we further assessed the number of AMR, virulence, and plasmid genes on a per isolate basis across all 42 countries for which we were able to obtain sequence data (Fig. 3). However, the distributions of AMR genes per isolate across all countries were still statistically significant in pairwise comparisons of the five AMR databases’ hit counts (P < 0.05). This demonstrated that the choice of reference database will still alter interpretation of AMR distributions across countries. This metric also provided a different list of the top 5 countries however, identifying countries with fewer than 10 isolates as carrying more AMR genes. Once filtered for countries with at least 10 isolates, only 26 countries were considered, of which the list of top five countries functional genes per isolate varied for each database. The top five countries with most genes per isolate for the argannot, NCBI AMRplus, and resfinder databases were the Democratic Republic of Congo, Malawi, Mali, Zambia, and Zimbabwe consistently, whereas for the megares database, the top five countries were Democratic Republic of Congo, Ghana, Mali, Zambia, and Zimbabwe, and for the card database the top five countries were Democratic Republic of Congo, Ghana, Kenya, Mali, and Zimbabwe. This highlights the likely variability when different AMR databases are used for data comparison. Democratic Republic of Congo, Mali, and Malawi were consistently in the top five countries across four AMR databases, and were also among the countries with the highest plasmids and virulence factors per isolate. However, these correlations are not adequate to predict origin or transfer of functional genes within or across countries.

**Figure 3.**
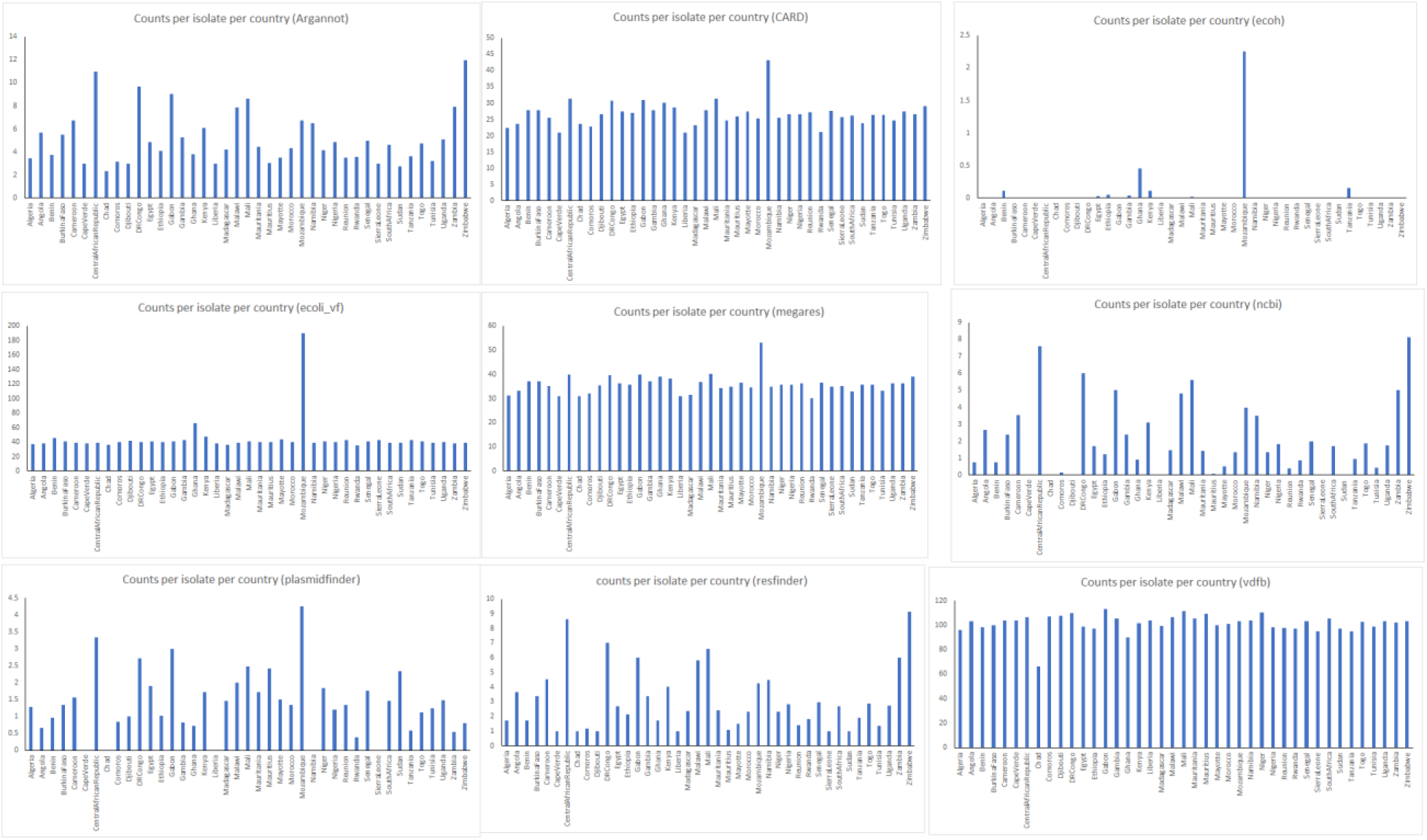
Country-wise distribution of various functional genes per isolate

Out of 4,552 isolates whose sequences were publicly available we were able to extract metadata from NCBI SRA for 1,002 samples that were isolated from various food sources. All these isolates had functional genes matches for each category. In order to understand the prevalence of *Salmonella* by food category in Africa, we applied the FAO/WHO’s food categorization scheme (Fig. 4 & 5) at level 2 categorization. We noted that the majority of food *Salmonella* isolates were derived from meat/poultry category (68%), within the terrestrial animals category (Fig. 6). However, eggs (0.33%) and dairy (1.66%) also contributed a minor fraction of *Salmonella* isolates from food from terrestrial animals. The next major contribution was within the plants category, where produce (15.9%) and nuts-seeds (9.5%) contained the most isolates vs grains-beans category (0.55%).

**Fig. 4.**
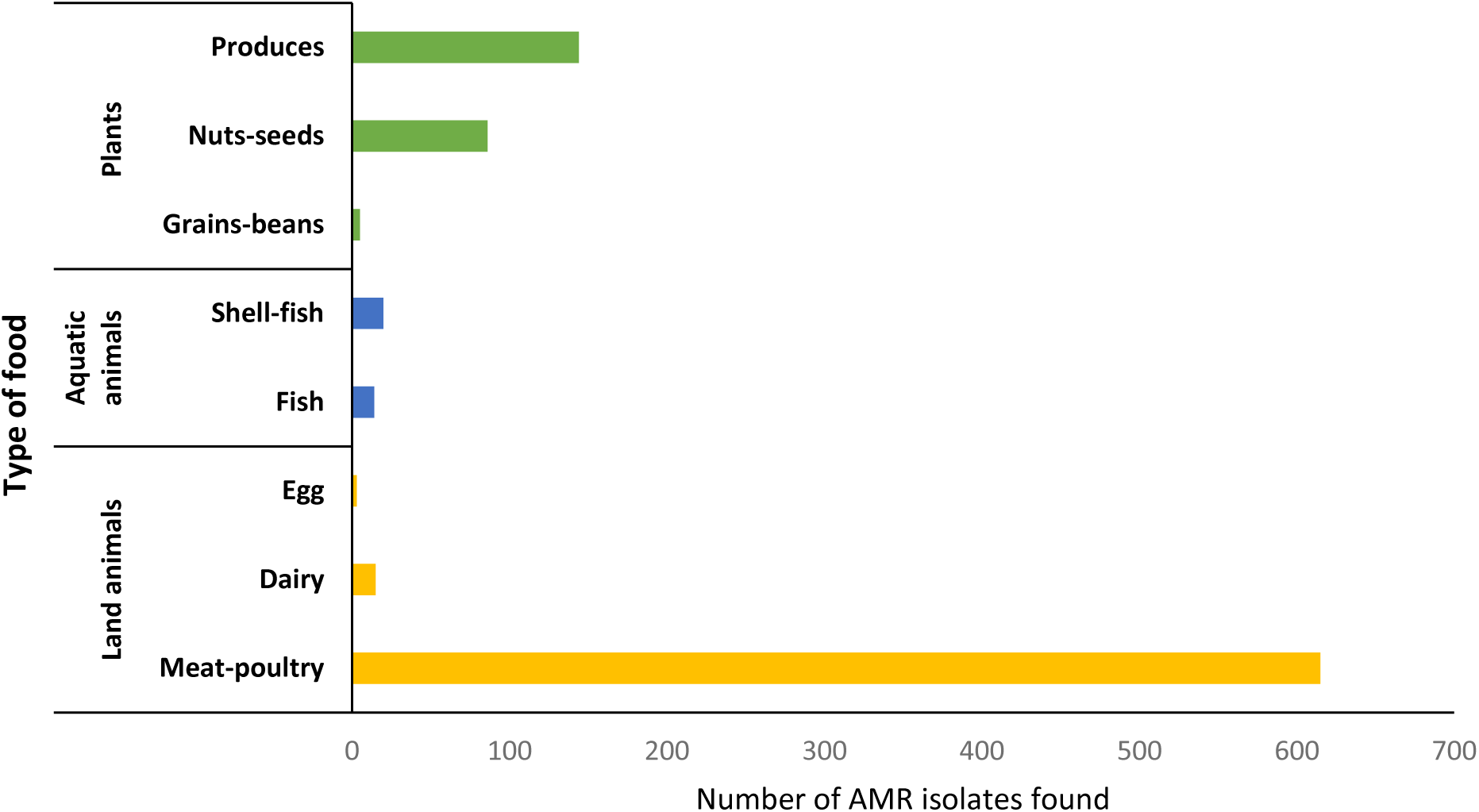
Isolates identifiable with a foodborne source

**Fig. 5.**
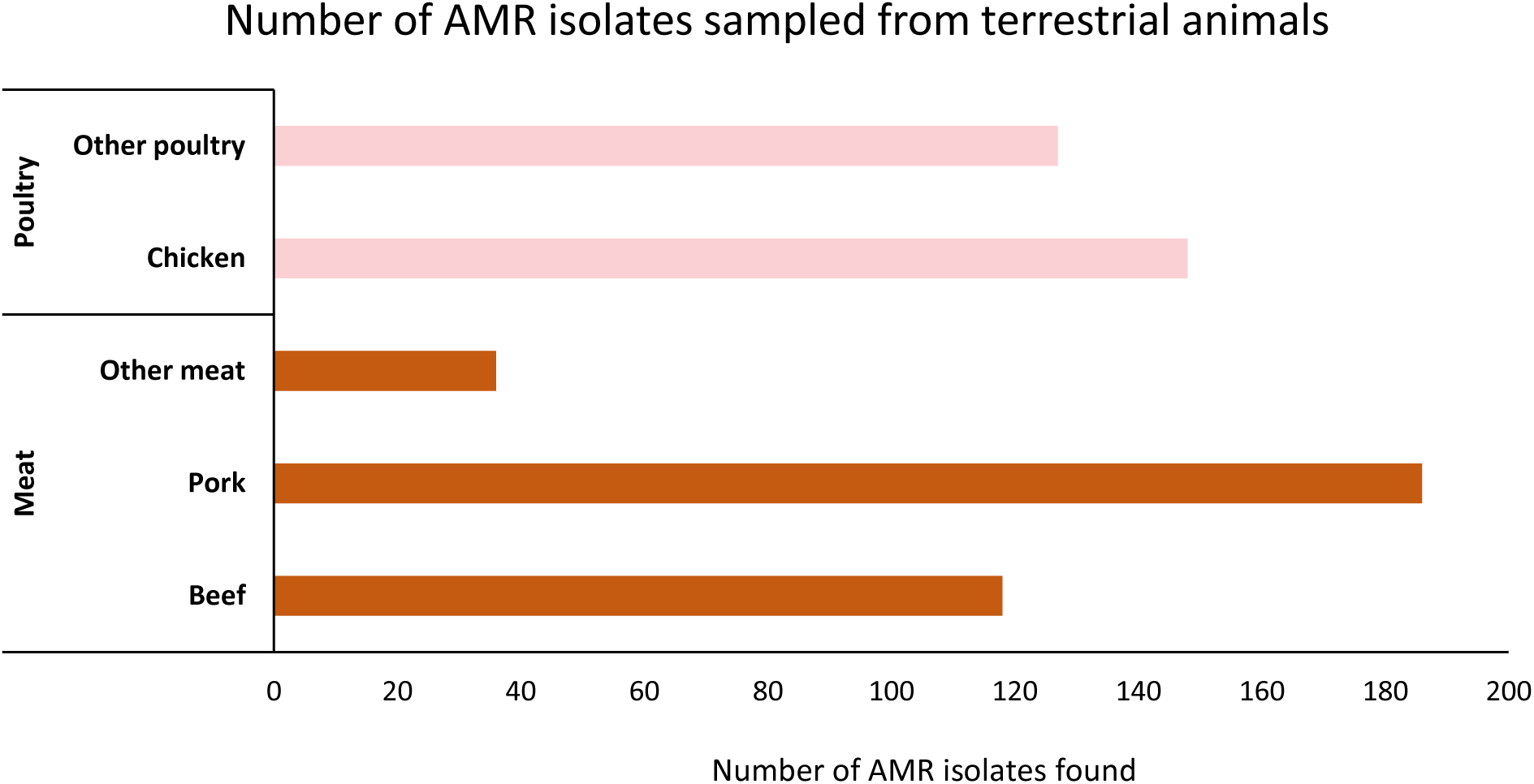
Isolates identifiable as from terrestrial animal sources

**Fig. 6.**
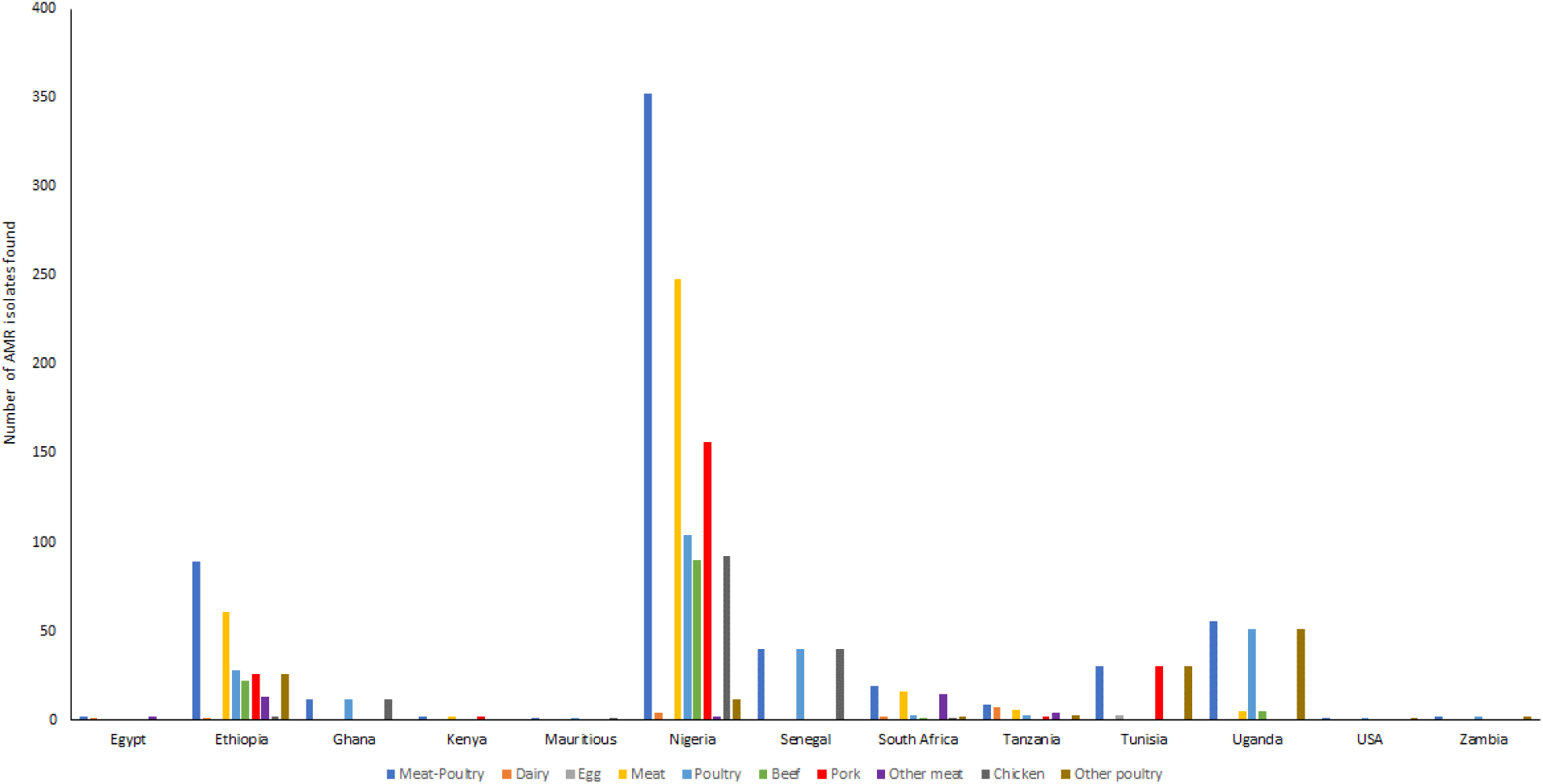
Country-wise distribution of *Salmonella* isolates from terrestrial animal sources

We further assessed country-wise food categories at levels 2, 3, and 4, which highlighted that the transmission mechanisms may vary across the African continent, due to variable food groups where AMR-bearing isolates were found. For example, AMR isolates from terrestrial animals were found only in 12 countries (Egypt, Ethiopia, Ghana, Kenya, Mauritius, Nigeria, Senegal, South Africa, Tunisia, Uganda, United Republic of Tanzania, and Zambia). In comparison, AMR isolates from aquatic animals were found in 9 countries (Gambia, Ghana, Kenya, Morocco, Nigeria, Sierre Leone, Tunisia, Uganda, and United Republic of Tanzania,) and from plant sources in 13 countries (Burkina Faso, Cameroon, Egypt, Ethiopia, Ghana, Kenya, Morocco, Nigeria, Rwanda, South Africa, Sudan Togo and Uganda). Notably, Ghana, Kenya, Nigeria, and Uganda have AMR bearing isolates from all food sources. Egypt and United Republic of Tanzania had AMR-bearing isolates in meat and dairy sources, whereas Morocco has AMR bearing isolates in shellfish and produce. This suggests that *Salmonella* in different countries may be found in different food niches and thus are likely to be genomically more diverse.

Genomic diversity may play a role in detected AMRs, thus, we further assessed the overall genomic diversity across all pairs of isolates. Notably, only Gambia and South Africa contained identical isolates, with 22 out of total 167 isolates forming 75 pairs that were identical in Gambia, and 16 isolates out of total 1,326 forming 8 pairs that were identical in South Africa. When comparing isolate pairs with fewer than 10 shared k-mers, within countries 585 isolate pairs were closely related, ∼0.03% of isolate pairs within the same country. Extending this range to fewer than 100 shared k-mers, 36,617 pairs were ancestrally related, ∼2.2% of all isolate pairs. We further performed similar comparisons across isolates not from the same country, where we found that no identical isolate pairs were found across 12,597,004 pairwise comparisons. Of these, 688 isolate pairs (∼0.005%) were found to share ≤ 10 k-mers and 35,344 isolate pairs (0.28%) shared > 10 k-mers and ≤ 100 k-mers. These comparisons highlight the low scale at which cross country transmissions are likely to occur at the African continent, based on our assessment of available sequence data till June 2021.

We further mapped the country locations of isolates and the number of pairs between countries where more than 10 isolate pairs were found to be closely related. Due to lack of geographical proximity among the related pairs (data not shown), and considering other studies showing that the transmission of *Salmonella* spp. and *Listeria* spp. across food processing facilities even within the same country is very low [28], these closely related pairs are unlikely to represent any cross border transmission potential within the African continent.

Expanding the shared k-mer range up to 100 showed that 22,535 pairs of isolates might be ancestrally related, though less likely to be recently transmitted. Of these, 3,564 isolate pairs were found to be in 17 pairs of countries that share borders, with the Malawi-Zambia country pair accounting for 1,665 pairs of isolates, effectively 46% of the total isolate pairs. Mali and United Republic of Tanzania shared isolates with 4 border countries, whereas Kenya shared isolates with 3 border countries. The rest of the isolate pairs across countries that don’t share borders are unlikely to be ancestrally related, despite genomic similarity. However, additional metadata beyond source country and deeper investigations would be needed to conclusively demonstrate cross border transmission of AMR gene bearing *Salmonella* isolates, beyond genomic comparisons alone.

Egypt and Ghana, the countries for which most isolates bearing AMRs were found in aquatic animals and plants categories did not feature among cross border related isolate country pairs even at 100 shared k-mers. Whereas Nigeria, for which the highest number of isolates were found in land animals, did share 399 pairs of isolates with its border neighbors Benin and Cameroon. This indicates that cross border isolate transmission may be region specific in the African continent and might only happen in limited instances. The number of AMRs found within these countries per isolate are very similar, for example 3.8-4.8 for argannot, 26.6-30.1 for card, and 35.8-39.1 for megares. Thus, the number of AMRs per isolate do not seem to influence cross border transmission. This is, however, challenging to predict effectively due to lack of any large-scale variation in AMRs per isolate across most countries (Fig. 3).

## 4. Discussion

We conducted this study to provide a broad understanding of AMR diversity in the African continent and to determine the impact of using specific AMR gene databases in identifying AMR genes. AMR is growing across the world and is considered one of the top challenges in microbial infectious disease prophylaxis. Surveys of AMR genetic potential in Africa are lacking. Here we chose *Salmonella* as the primary genus to help us understand foodborne prevalence of AMR genes.

We found that under relatively stringent parameters for nucleotide identity, ≥ 75% of matches were identical to known functional genes, and ≥ 99% matches were 90% genetically identical. This suggests that ∼25% genetic variation is prevalent in certain functional gene categories. We accept that this genetic diversity is to be expected. This could mean that certain genes may need to be more conservatively analyzed for their ability to be functional. Such analysis was not conceived as part of the present genomic survey of AMRs in *Salmonella* and would be beneficial for future perspectives.

Recently, comparable studies have identified among limited numbers of *Salmonella* isolates the genetic potential of AMR & other functional genes at Burkina Faso [29], Kenya [30,31], Malawi [32], Nigeria [33], and United Republic of Tanzania [34]. Increase in AMR has also been identified via phenotypic surveillance methods on some African countries such as Egypt [35,36], Ethiopia [37,38], South Africa [39], and Uganda [40]. Wildlife in Africa also carry *Salmonella* that harbor AMR genes [41]. Additionally, AMR prevalence continent-wide has been documented by reviews and meta analyses [40,42–44]. Here, we surveyed the *Salmonella* isolates available across the African continent to identify whether there is AMR prevalence and to help us understand whether AMR prevalence estimates are likely influenced by the choice of genetic databases for these functional genes.

The curated databases against which bacterial genomes are compared for AMR potential appear to be a key aspect, as they may provide a significantly variable picture. For example, we found that across 5 well known AMR gene databases, the average numbers of AMR genes may vary from as low as 5 to as many 28 (Fig. 2). Also, the number of *Salmonella* isolates that carry AMR genes may vary depending on the choice of the database. Such diverse interpretations would limit our ability to understand and interpret AMR predictions. For prominent use in AMR surveys, harmonized standards need to be accomplished across various curated databases of functional genes. Scientific efforts and contributions by various teams in creating these high quality databases will be empowered for better utilization with globally harmonized standards that integrate the contents of all these databases.

One aspect that is recently being commonly adapted is the extensive use of metadata in genomic surveys to allow us dive into various contexts [45]. FDA in partnership with other US-based agencies has conducted significant efforts to create a specific taxonomy for metadata (https://www.fda.gov/science-research/fda-science-forum/standardizing-isolation-source-metadata-genomic-epidemiology-foodborne-pathogens-using-lexmapr), using a tool called LexMapr (https://github.com/cidgoh/LexMapr). At the time of conduct of this study, such metadata taxonomy schemes were not available.

We supported our analyses in this situation by adopting the FAO’s food category taxonomy to the 1,002 isolates where food source information was available, 22% of total isolates we were able to obtain from INSDC. This subgroup analysis however did help to identify that various local food preferences may likely modify the possibility of cross border transmission of AMR bearing *Salmonella* inside the African continent. Other isolates not originating from food related sources and most likely originating from clinical samples, we chose to look at overall genomic diversity outside of the food categorization scheme to confidently assess cross border transmission of *Salmonella* inside Africa.

We assessed the geographical distribution of genomic diversity and the likelihood that AMR bearing *Salmonella* may frequently distribute across geographic borders. Our assessment indicates that such cross-border transmission is likely to be limited to certain endemic regions where fewer than 100 k-mers were distinct between countries sharing borders. Other studies have shown that even at fewer than 10 SNP differences, sharing of *Salmonella* isolates is likely to be very low even across food processing facilities that are not directly linked to each other [28].

Taking this observation and our assessments in context, isolates that differ by ≤ 10 or ≤ 100 shared k-mers across countries that don’t share borders are not likely to be indicative of cross border *Salmonella* or AMR transmission.

K-mer distribution is prevalently used to address genome scale diversity for taxonomic identification and phylogeny at species & subspecies level [46–49], and is not frequently applied for tracing back to source of origin. We have used k-mer analysis in our study to demonstrate that for larger scale surveys to assess genome diversity, k-mer analysis can be quite powerful and facile for data interpretation. Mainly however, SNP-based analyses are more resource intensive [50], and are thus do not scale to comparisons of 100’s of isolates. In some instances, k-mer analyses are used to limit prior genomic diversity for SNP-based analyses [28,51]. Such an approach can be both useful and facile for interpreting smaller branches of limited genomic diversity for more focused source tracking analyses. International Organization for Standardization (ISO) guidelines from 2020-2021 for genomic analysis of foodborne bacteria do not recommend k-mer-based analyses for source tracking, restricting user choice recommendations to SNP or genomic scale MLST approaches (ISO 16140-3:2021). More recently, WHO has also issued guidelines for foodborne bacterial analyses and source tracking in 2023 (https://www.who.int/publications/i/item/9789240021228) where the use of k-mer analyses is equally recognized alongside SNP and genomic MLST. These observations only indicate that the field of bacterial source tracking is scientifically evolving and a consensus is yet to be established. Further assessment and use of k-mer analyses is certainly suited to broad scale surveys where interpretation may happen within minutes as opposed to likely much longer operational times for SNP and genomic MLST pipelines. K-mer analysis needs to be critically evaluated in cohort with SNP & genomic MLST in light of emerging science with high quality benchmark data to validate the tool’s applicability in real life source tracking and genetic potential exchange scenarios.

## 5. Conclusions

Antimicrobial resistance genes, functional genes for virulence, and plasmid-borne genes were identified in 4,552 *Salmonella* isolates by assembling genomes from publicly available unassembled data. The choice of database may alter interpretation of AMR prevalence. Harmonizing the genetic content of such functional gene databases will lend to uniformity in global assessment of AMR gene transmission. Meat/poultry was the chief food category for foodborne *Salmonella*. Based on genome comparisons, cross border transmission of *Salmonella* isolates happens at very low frequency and may be related to food preferences.

---

**Supplementary data Figure S1.**
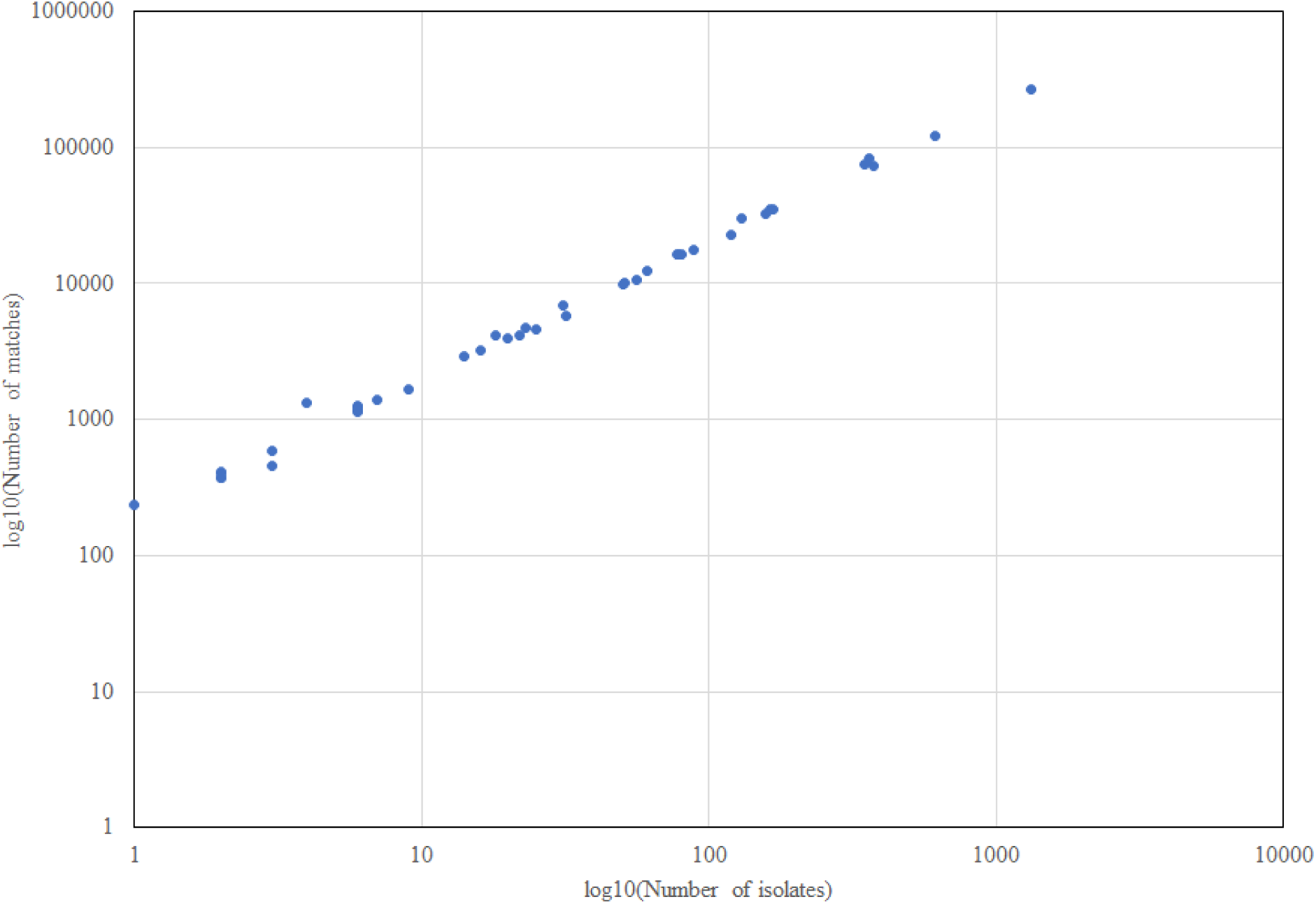
Number of AMR genes, virulence factors and plasmid-borne genes carried by all isolates in African countries

